# A Combination Strategy of Solubility Enhancers for Effective Production of Soluble and Bioactive Human Enterokinase

**DOI:** 10.1101/2021.07.22.453328

**Authors:** Jinhak Kwon, Hyeongjun Cho, Seungmin Kim, Yiseul Ryu, Joong-jae Lee

## Abstract

Enterokinase is one of the hydrolases that catalyze hydrolysis to regulate biological processes in intestinal visceral mucosa. Enterokinase plays an essential role in accelerating the process of protein digestion as it converts trypsinogen into active trypsin by accurately recognizing and cleaving a specific peptide sequence, (Asp)4-Lys. Due to its exceptional substrate specificity, enterokinase is widely used as a versatile molecular tool in various bioprocessing, especially in removing fusion tags from recombinant proteins. Despite its biotechnological importance, mass production of soluble enterokinase in bacteria still remains an unsolved challenge. Here, we present an effective production strategy of human enterokinase using tandemly linked solubility enhancers consisting of thioredoxin, phosphoglycerate kinase or maltose-binding protein. The resulting enterokinases exhibited significantly enhanced solubility and bacterial expression level while retaining enzymatic activity, which demonstrates that combinatorial design of fusion proteins has the potential to provide an efficient way to produce recombinant proteins in bacteria.

## 1. Introduction

Along with optimization of biological activity, soluble expression of recombinant active proteins is one of the major issues to be considered in the field of biotechnology, in terms of cost-effectiveness and mass production of biological reagents. However, some valuable proteins, mostly derived from human and other eukaryotes, are prone to form insoluble and inactive aggregates while using a bacterial expression system due to improper protein folding and modification (Butt, Edavettal, Hall, & Mattern, 2005; Pacheco, Crombet, Loppnau, & Cossar, 2012). To address this challenge, several advanced strategies have been introduced to achieve effective and stable expression of soluble recombinant proteins (Khow & Suntrarachun, 2012). Among them, the use of solubilizing fusion partners including maltose-binding proteins (MBP) (Dyson, Shadbolt, Vincent, Perera, & McCafferty, 2004; Kapust & Waugh, 1999; Song, Lee, Park, Han, & Lee, 2012) and glutathione S-transferase (GST) (Rabhi-Essafi, Sadok, Khalaf, & Fathallah, 2007) has been recognized as the primary option for improving protein solubility and expression level. In addition, fusion tags can be utilized for facile purification of expressed proteins by affinity chromatography, allowing high-yield purification with remarkable purity (Waugh, 2005). Despite these advantages, the removal of the fusion partners from purified proteins is highly recommended for further applications because they can inhibit the biological function of recombinant proteins and trigger unwanted immune responses when used as a therapeutic agent (Choi, Song, Moon, & Seong, 2001). For this process, a variety of endopeptidases have been commonly used for cleavage of pre-designed peptide linkers located between the solubilizing protein and the protein-of-interest (Waugh, 2011).

Hydrolases are a group of hydrolytic enzymes that promote the breakdown of various biochemical bonds. Among them, protease (peptidase) has a specialized role in cleavage of amide bonds in peptides and proteins to precisely regulate biological processes and to maintain homeostasis (Mahesh, Tang, & Raj, 2018). Enterokinase, also known as enteropeptidase, belongs to a class of serine proteases and is a disulfide-linked heterodimeric enzyme, composed of a heavy chain (87 kDa) and a light chain (26 kDa) (Kitamoto, Yuan, Wu, McCourt, & Sadler, 1994). In the small intestine, enterokinase is mainly produced in the duodenal brush borders for activation of the digestive system by converting proenzyme trypsinogen into active proteolytic enzyme, trypsin (Lobley et al., 1973; Yamashina, 1956). Interestingly, the light chain of enterokinase exhibits a remarkable target specificity and catalytic activity, enabling the effective recognition of a pentapeptide sequence Asp-Asp-Asp-Asp-Lys (DDDDK) and hydrolysis of the bond located after the lysine residue (Light & Janska, 1989). With this site-specific activity, enterokinase is found to be stable over a broad range of temperature (4∼45 °C) and pH (4.5∼9.5) (Gasparian, Ostapchenko, Dolgikh, & Kirpichnikov, 2006). Based on these properties, enterokinase has been widely applied as a key enzyme for the process of isolating and removing the solubilizing fusion partners or affinity tags from purified recombinant proteins (Choi, Song, Moon, & Seong, 2001). However, several difficulties exist in enterokinase production. Previous studies reported that overexpressed enterokinase in bacteria was dominantly expressed in insoluble aggregates and formed inclusion bodies, resulting in extremely low production yields of soluble enterokinase (Gasparian, Ostapchenko, Dolgikh, & Kirpichnikov, 2006; Niu, Li, Ji, & Yang, 2015). In order to address this limitation, a refolding method was introduced and implemented despite the apparent disadvantage of a complicated and inconvenient process (Yi & Zhang, 2006; Tan, Wang, & Zhao, 2007). In addition, periplasmic expression of enterokinase was attempted but showed low expression levels (Collins-Racie et al., 1995). Due to the difficulty of obtaining recombinant enterokinase in bacteria, most of the commercial enterokinases are expressed in yeast or extracted from the intestine of animals with high production costs (Melicherová et al., 2017).

This study presents a soluble expression of recombinant enterokinase light chain (EKL) in *Escherichia coli* (*E. coli*) by using several solubility-enhancing tags (**Scheme 1**). As human enterokinase is known to have 10 times higher catalytic activity than animal enterokinase, human enterokinase is selected for this study and further applications (Gasparian et al., 2006). As shown in **Figure S1**, unmodified enterokinases exhibit poor solubility and low expression level in *E. coli*. To increase solubility of expressed enterokinases, we employed two types of fusion proteins including maltose binding protein (MBP) (di Guan, Li, Riggs, & Inouye, 1988) and phosphoglycerate kinase (PGK) (Park et al., 2008; Song, Lee, Park, Han, & Lee, 2012). Thioredoxin (Trx) was also used to facilitate correct disulfide-bond formation and protein folding, as enterokinase contains eight cysteine residues (Nordström, 1972). These protein tags were genetically incorporated into the N-terminus of enterokinases in various combination. The resulting enterokinases were highly expressed in *E. coli* as a soluble form with proteolytic activity comparable to a commercial protein, as the details demonstrated herein.

**Scheme 1.**
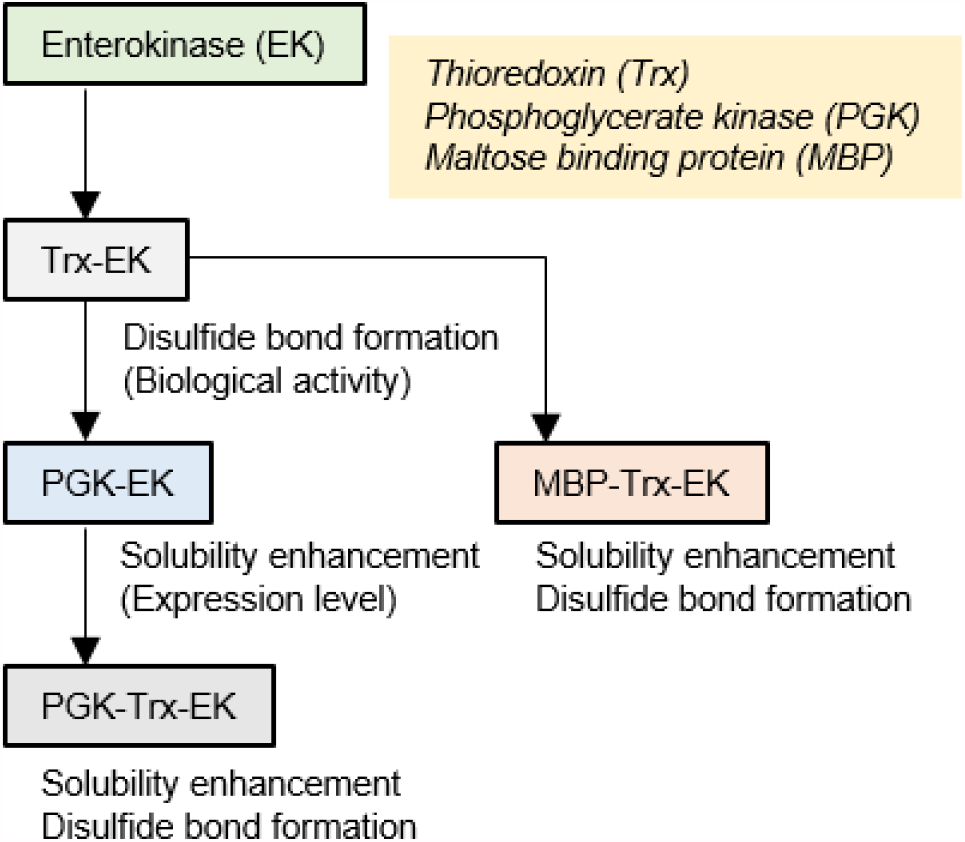
Schematic representation of a combinatorial approach of solubilizing proteins for effective bacterial expression of enterokinase. Each fusion protein was genetically introduced based on their intrinsic biochemical properties. Thioredoxin was selected for promoting disulfide bond formation and proper protein folding. Maltose-binding protein and Phosphoglycerate kinase were employed for solubility enhancement of their fusion partner, enterokinase.

## 2. Materials and Methods

### 2.1. Construction of expression vectors for enterokinase protein production

The gene coding for human enterokinase light chain (EKL) was synthesized (IDTdna, USA), and amplified through PCR using a forward primer bearing *NdeI* cleavage site and a reverse one with *XhoI*. For appropriate folding and enhanced solubility, thioredoxin (Trx) or phosphoglycerate kinase (PGK) gene was additionally incorporated before hEKL (Trx-hEKL and PGK-hEKL) by overlap extension PCR. A linker between Trx and hEKL is the same as the one used in a pET-32a (+) vector (Novagen, USA) except a mutation of *NdeI* cleavage site and removal of His-tag in the linker. PGK-hEKL has a 15-amino acid linker between the two proteins. Furthermore, to confer both proper disulfide-bond formation and high solubility, maltose binding protein (MBP) or PGK gene was linked to Trx-hEKL (MBP-Trx-hEKL and PGK-Trx-hEKL) with a 10-amino acid linker by overlap extension PCR using each solubilizing tag and Trx-hEKL as a PCR template. All constructs except PGK-hEKL possessed one cleavage site by enterokinase (DDDDK) right before hEKL. These four types of differently tagged hEKL were amplified finally through PCR using a forward primer bearing *NdeI* cleavage site and a reverse one with *XhoI*. The resulting five constructs were cloned into a pET-21a (+) vector (Novagen, USA) between the *NdeI* and *XhoI* sites. **Table S1** presents all complete amino acid sequences constructed in this study.

### 2.2. Expression and purification of enterokinases

All types of hEKL-containing pET21a plasmid vectors were transformed into Origami B (DE3) competent cells, and the transformed cells were incubated on LB broth agar plates containing ampicillin (100 μg/ml), kanamycin (50 μg/ml), and tetracycline (10 μg/ml) overnight at 37°C. A single colony on each construct was inoculated into a 10 ml LB broth medium and cultured at 37°C overnight. The culture broth was diluted 100-fold in a 1 L LB medium containing three types of antibiotics and incubated further at 37°C in a shaking incubator until optical density at 600 nm (OD_600_) reached 0.6-0.8. When the OD_600_ reached that point, Isopropyl-ß-D-thiogalactopyranoside (IPTG) was added at final concentration of 0.1 mM to induce protein expression. The cells were further grown at 18°C overnight. Cell harvest was conducted by using centrifugation at 4000 rpm for 30 min at 4°C The pelleted cells were re-suspended with a 45 mL lysis buffer (20 mM Tris-Cl (pH 8.0), 300 mM NaCl, and 10 mM Imidazole) and disrupted by sonication. The lysate was centrifuged at 13,000 rpm for 1 h at 4°C to save the supernatant. The soluble hEKL proteins were purified using a Ni-NTA agarose resin (Qiagen, Germany) and subsequently HiLoad 16/60 superdex 75 prep grade column (GE Healthcare, USA). The purity of each hEKL was estimated by SDS-PAGE.

### 2.3. Enzyme cleavage assay

To detect the cleavage activity of the recombinant hEKL constructs, Trx-tagged repebody (Trx-rH9A4) harboring the enterokinase recognition sequence DDDDK in the linker was used as a substrate protein. This substrate protein was prepared according to the procedure described above using rH9A4 gene-inserted pET32a vector. All the proteins used in a cleavage assay were buffer-exchanged with an enterokinase reaction buffer (20 mM Tris-HCl (pH 8.0), 50 mM NaCl and 2 mM CaCl_2_) by PD-10 desalting column (GE Healthcare, USA). The cleavage reaction was performed with hEKL (370 nM) and Trx-rH9A4 (1 mg/ml, 12.3 μM) in an enterokinase reaction buffer at 37°C overnight. To analyze relative activity of the recombinant hEKL, commercial bovine enterokinase / PRSS67 protein (Sino Biological, China) was used as a positive control. The cleavage activities of each hEKL were determined by SDS-PAGE analysis.

### 2.4. Measurement of bacterial expression level of enterokinases

To evaluate the expression level of soluble hEKL proteins, soluble fraction of each recombinant hEKL protein after cell lysis was diluted to detect a clear protein band through SDS-PAGE. For quantification of the hEKL proteins, known concentrations of BSA (Thermo Scientific, USA) proteins (1.0 mg/ml, 0.5 mg/ml, and 0.25 mg/ml) were used as standards, and electrophoresis of all the protein samples was conducted. After SDS-PAGE, the gels were analyzed with Image J program (National Institutes of Health, USA), and the concentrations of enterokinases in the diluted soluble fractions were calculated using BSA standard curve. The expression level (mg/L) was determined by multiplying the concentration by the lysis volume of liter culture.

### 2.5. Determination of solubility of enterokinases

The enterokinases located in soluble and insoluble fractions were analyzed through SDS-PAGE. For preparing the insoluble fraction, a centrifuged pellet after cell lysis was dissolved with 8 M urea solution in the same volume as a lysis buffer. Enterokinase protein bands in the soluble and insoluble fractions were quantified through Image J program, and the relative solubility was determined as follows: Band intensity in a soluble fraction / (BI in a soluble fraction + BI in an insoluble fraction) x 100%.

## 3. Results and Discussion

### 3.1. Bacterial expression of human enterokinase fused with a thioredoxin tag

Human enterokinase (EK) consists of two distinct domains, the heavy and light chains. Of the two, the light chain (26.5 kDa) is known as the catalytic domain responsible for enzymatic cleavage of specific peptide sequences. In this research, the cysteine-rich light chain of enterokinase was selected to conduct soluble expression in a bacterial system (**Figure S2**). First, the catalytic domain of EK protein was genetically fused with thioredoxin (Trx) to promote the proper formation of disulfide bonds, which substantially affect protein stability and activity. (**Figure 1a**). With this, the target amino acid sequence (DDDDK) for enterokinase was inserted between Trx tag and EK protein to obtain the intact form of enterokinase by autocleavage. The resulting Trx-fused enterokinase light chain (Trx-EKL) was expressed in *E. coli* strain Origami B (DE3). As demonstrated in **Figure 1b**, SDS-PAGE analysis displayed an induced band of Trx-EKL proteins in the soluble fraction although the majority of the proteins were shown to be insoluble as previously reported (Gasparian, Ostapchenko, Schulga, Dolgikh, & Kirpichnikov, 2003). Interestingly, Trx-EKL proteins underwent autocleavage after affinity purification, which seems to be the effect of removing the non-specific binding between *E. coli*-derived proteins and enterokinase during the purification process. To increase protein purity, size-exclusion chromatography (SEC) was subsequently performed, and the catalytic subunit of enterokinase with high purity (∼90%) was finally obtained (**Figure 1c**). Next, to verify whether purified enterokinases were biologically active, Trx-repebody fusion proteins (Trx-Rb) bearing DDDDK sequence were employed as a substrate for enterokinase cleavage assay. As a result, the resulting EKL proteins appear to have site-specific protease activity similar to commercially available ones (**Figure 1d**). These results demonstrate that the thioredoxin fusion tag allows low-level expression of poorly soluble EKL proteins in bacteria while retaining full enzyme activity.

**Figure 1.**
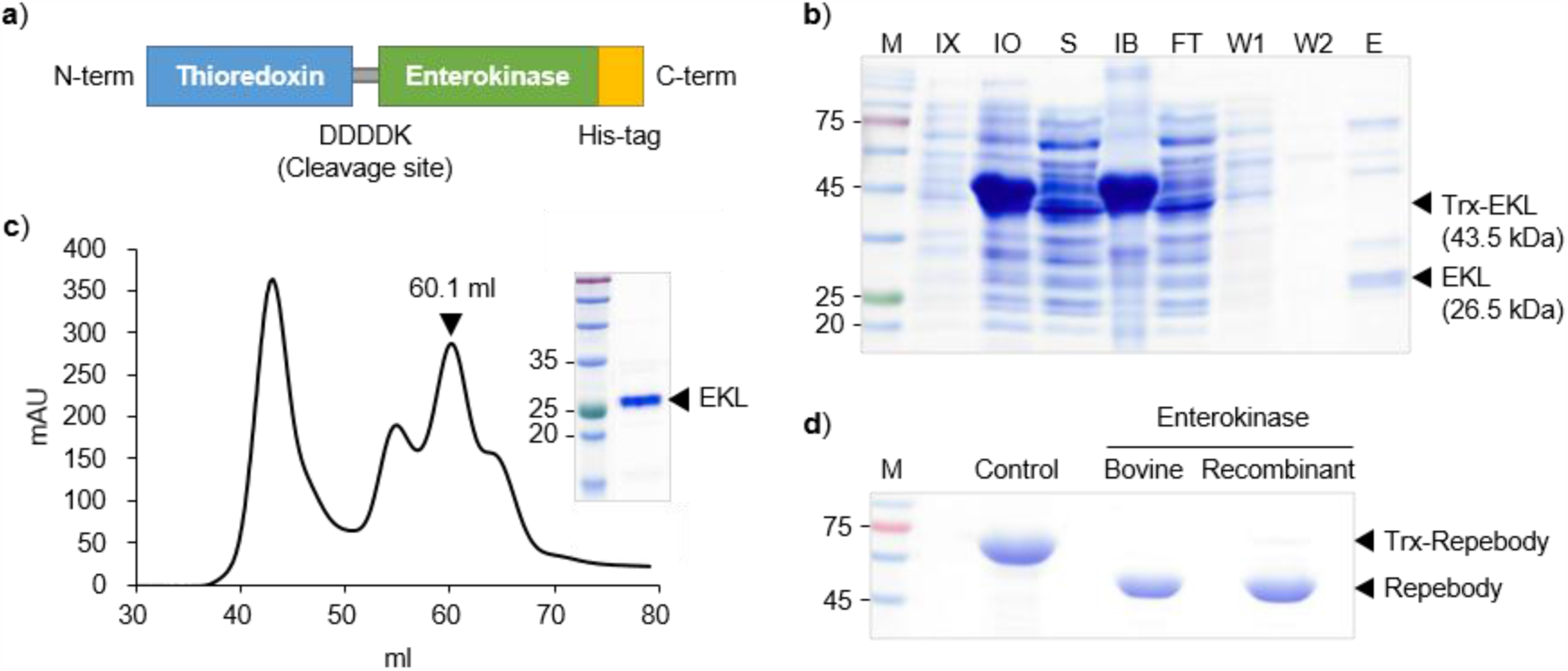
Expression and purification of thioredoxin-tagged enterokinase. **a**) Human enterokinase light chain (EKL) was genetically fused with thioredoxin (Trx) tag. DDDDK sequence was incorporated to separate EKL from Trx proteins after autocleavage. **b**) SDS-PAGE analysis of expressed Trx-EKL protein in bacteria. Expressed protein was purified by using nickel-nitrilotriacetic acid (Ni-NTA) resin. IX: Before IPTG induction. IO: After IPTG induction. S: Soluble fraction. IB: Inclusion body. FT: Flow through. W: Washing fraction. E: Elution fraction. **c**) Size-exclusion chromatography profile of purified EKL protein. Highly purified EKL was verified by SDS-PAGE analysis. **d**) Protein cleavage assay for identifying the proteolytic activity of the resulting EKL compared to commercially available bovine enterokinase. Trx-fused repebody was employed as a substrate for EKL, and DDDDK site was placed internally between Trx tag and repebody.

### 3.2. Improvement of soluble expression of human enterokinase using phosphoglycerate kinase

In biotechnology, achieving high production yields of recombinant proteins is a fundamental requirement for the successful development of biological products, which is as important as maintaining their intrinsic bioactivity. Unfortunately, thioredoxin-fused enterokinase (Trx-EKL) showed a very low-level expression in *E*.*coli*. To improve production yields, *E*.*coli* phosphoglycerate kinase (PGK) was introduced and genetically fused to the N-terminal end of enterokinase (**Figure 2a**). The PGK protein is one of the ten enzymes involved in glycolysis pathway, which participates in the conversion of 1,3-bisphosphoglycerate to 3-phosphoglycerate (Rojas-Pirela et al., 2020). More importantly, like other glycolytic enzymes, the PGK protein is constitutively expressed at high levels in bacteria (Liebermeister et al., 2014). Based on this property, previous studies have indicated that the PGK protein has a potential to be used as a fusion partner to increase the solubility of poorly expressed proteins in *E*.*coli* (Song et al., 2012). We expressed the PGK-fused enterokinase (PGK-EKL) in Origami B host cells. As expected, the solubility of PGK-EKL proteins was dramatically enhanced compared to Trx-EKL (**Figure 2b** and **Table 1**). However, most of the soluble proteins did not bind to the Ni-NTA affinity column, and they were washed out with binding buffer. The results suggest that the PGK protein does not contribute significantly to proper protein folding, so the C-terminal his-tag may be buried in the improperly folded protein, resulting in poor immobilization of PGK-EKL on the Ni-NTA resin. After washing, bound PGK-EKL proteins were eluted and subsequently introduced to SEC for further purification. Consequently, the purified PGK-EKL appeared to have been eluted as a single and narrow peak with high purity (∼95%) (**Figure 2c**). The finally purified PGK-EKL was subjected to the protein cleavage assay. As shown in **Figure 2d**, prominent amounts of the substrate protein as a full form were detected after the cleavage reaction with PGK-EKL, whereas nearly all proteins were cleaved by Trx-EKL. Considering the importance of disulfide bonds in protein folding and structural integrity, the reduced functionality of the PGK-EKL protein clearly indicated that PGK has limitations as a fusion partner in producing disulfide bond-rich proteins.

**Figure 2.**
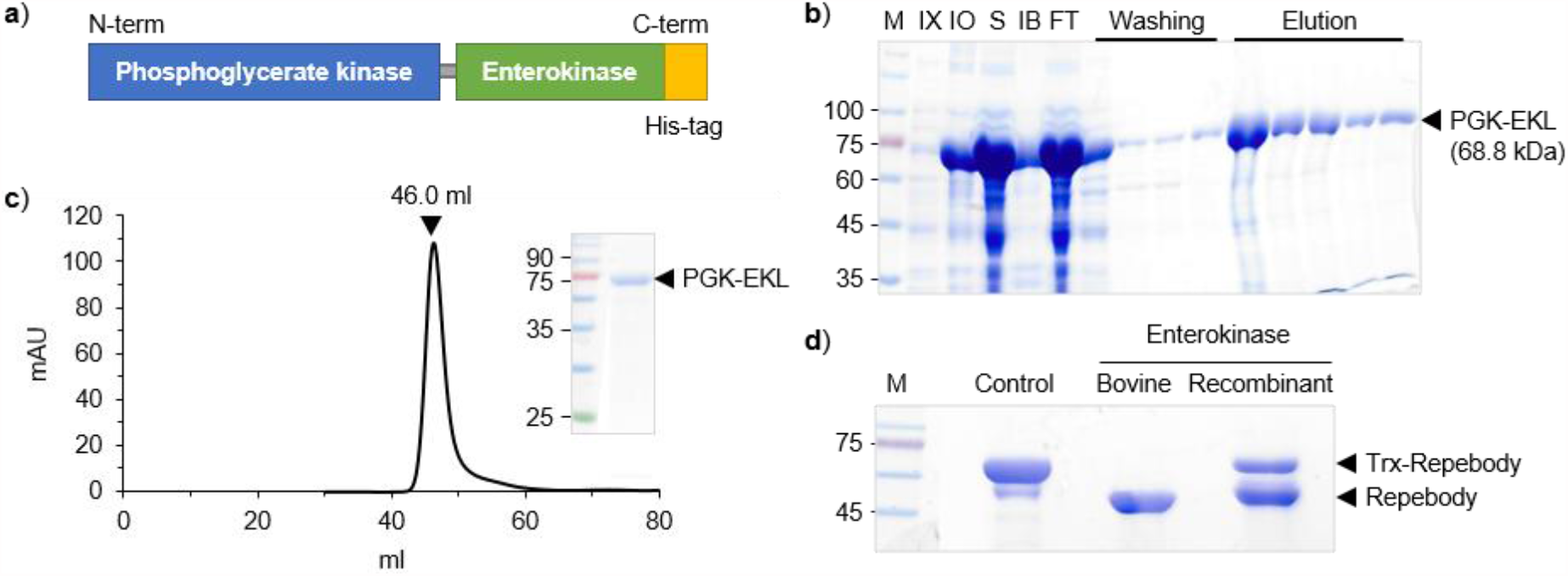
Expression and purification of phosphoglycerate kinase-tagged enterokinase. **a**) Schematic of phosphoglycerate kinase (PGK)-fused enterokinase (EKL). **b**) Bacterial expression of PGK-EKL protein. After affinity chromatography, purified proteins were visualized by SDS-PAGE analysis. The resulting PGK-EKL exhibited a significantly increased solubility compared to Trx-tagged EKL. **c**) Size-exclusion chromatography profile of purified PGK-EKL protein. PGK-EKL was eluted as a single peak with high purity. **d**) Protein cleavage assay showed that the enzymatic activity of PGK-EKL protein was partially reduced.

**Table 1.** Biochemical properties of engineered enterokinases in this study

| **Protein** | **Size (kDa)** | **Expression level**<br>**(mg/L culture)** | **Solubility (%)** |
| --- | --- | --- | --- |
| EKL | 26.5 | N.D | N.D |
| Trx-EKL | 43.5 | N.D | N.D |
| PGK-EKL | 68.8 | 101.33 | 92.5 |
| MBP-Trx-EKL | 84.6 | 142.20 | 94.5 |
| PGK-Trx-EKL | 84.7 | 175.95 | 95.0 |

EKL: Enterokinase light chain, Trx: Thioredoxin, PGK: Phosphoglycerate kinase, MBP: Maltose-binding protein, Solubility of each protein was calculated by comparison of relative band intensities of SDS-PAGE for soluble fraction (S) and inclusion body (IB).

### 3.3. Genetic incorporation of dual solubility-enhancing tags for optimized expression of functional human enterokinase

Considering the results presented above, the formation of disulfide bonds is of great importance as well as enhancement of solubility for effective expression of bioactive enterokinase in bacteria. In case of thioredoxin, although it did not substantially increase the soluble expression of EKL, the resulting proteins were shown to have fully retained biological activity. We thus constructed two additional types of EKL proteins based on Trx-fused form (**Scheme 1**). Maltose-binding protein (MBP) is well known as one of the most useful fusion tags for increasing protein solubility. However, previous studies reported that although MBP fusion can promote soluble expression of EKL as seen with PGK, it lacks the ability to form disulfide bonds properly (Kim, Lee, Park, Kim & Ahn, 2021). Therefore, we hypothesized that by taking advantage of each protein through simple genetic fusion, it is possible to effectively produce structurally and functionally folded enterokinase. Based on this concept, each solubility enhancer (MBP or PGK) was genetically introduced at the N-terminal end of Trx-EKL to achieve high-level soluble expression of bioactive enterokinase (**Figure 3a** and **Figure 4a**). As presented in **Figure 3b**, MBP-tagged Trx-EKL has remarkably enhanced solubility with the negligible formation of inclusion body. After purification, the proteolytic ability of MBP-Trx-EKL was verified through self-cleavage assay (**Figure 3c**). Under two different temperature conditions, MBP-Trx-EKL underwent effective self-cleavage at room temperature (RT), but the cleaved EKL was highly aggregated at 4^º^C. Therefore, further purification steps were carried out using size-exclusion chromatography (SEC) at RT. The self-cleaved MBP-Trx-EKL was separated into two distinct peaks, and the cleaved EKL was observed in an earlier eluting peak (Fraction A) (**Figure 3d**). The resulting EKL effectively cleaved all given substrate proteins (**Figure 3e**), which is similar to Trx-EKL. These results imply that a cooperative action of the two fusion proteins (MBP and Trx) can lead to high-level soluble expression of biologically active human enterokinase in bacteria.

**Figure 3.**
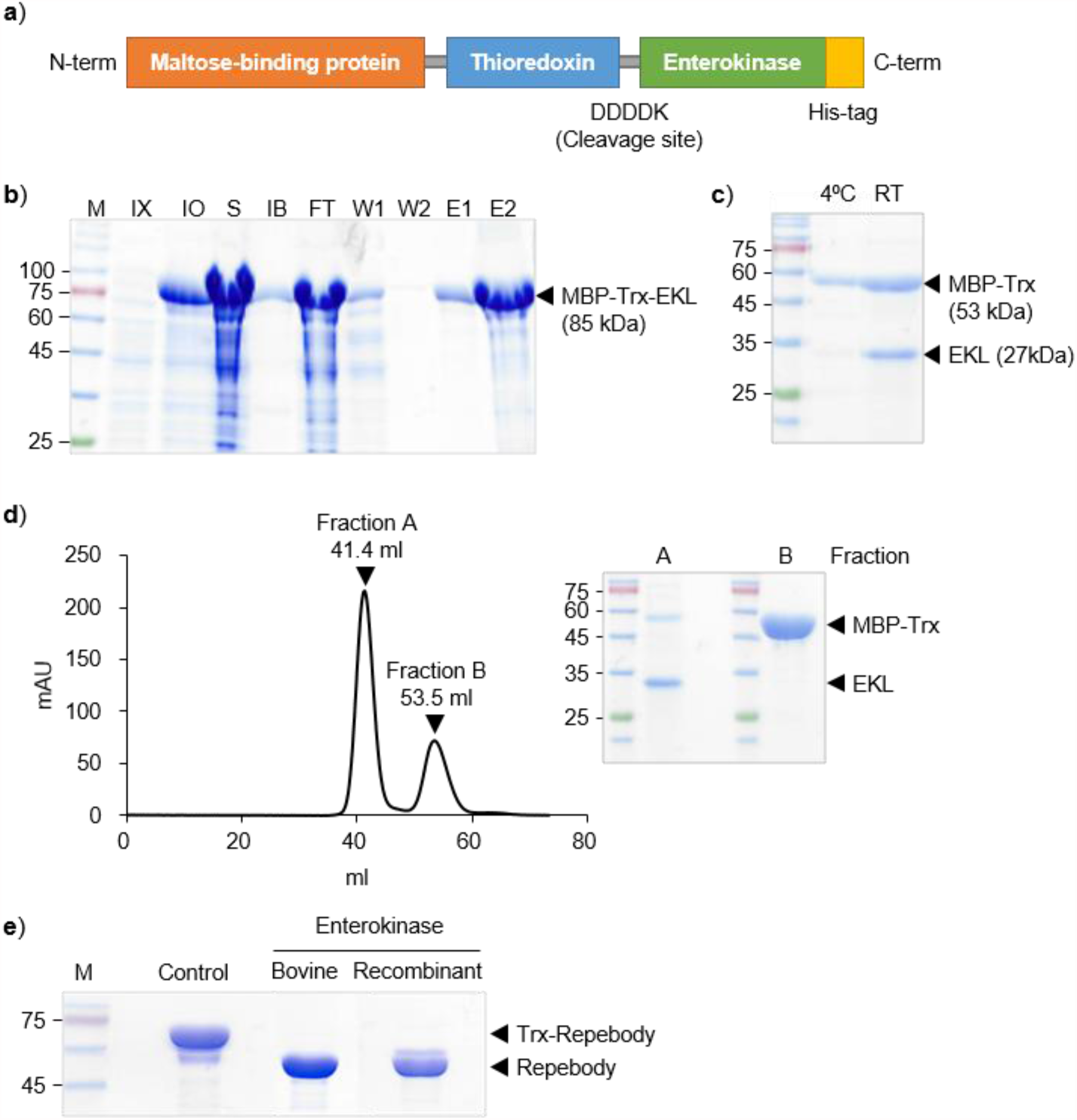
Improvement of solubility of thioredoxin-tagged enterokinase (Trx-EKL) through genetic incorporation of maltose-binding protein. **a**) To increase soluble expression level of Trx-EKL, maltose-binding protein (MBP) was fused at the N-terminal end of Trx-EKL protein. **b**) Remarkably enhanced solubility and expression level of MBP-tagged Trx-EKL were confirmed by SDS-PAGE analysis. **c**) Optimization of the autocleavage process of MBP-Trx-EKL. Cleaved EKL protein can be highly obtained at room temperature. **d**) Size-exclusion chromatography profile of MBP-Trx-EKL protein. After cleavage reaction, MBP-Trx-EKL was subsequently introduced to chromatography and free EKL was separated from digested MBP-Trx. **e**) Protein cleavage assay revealed that MBP-Trx-EKL has similar biological activity to parent Trx-EKL with high solubility.

**Figure 4.**
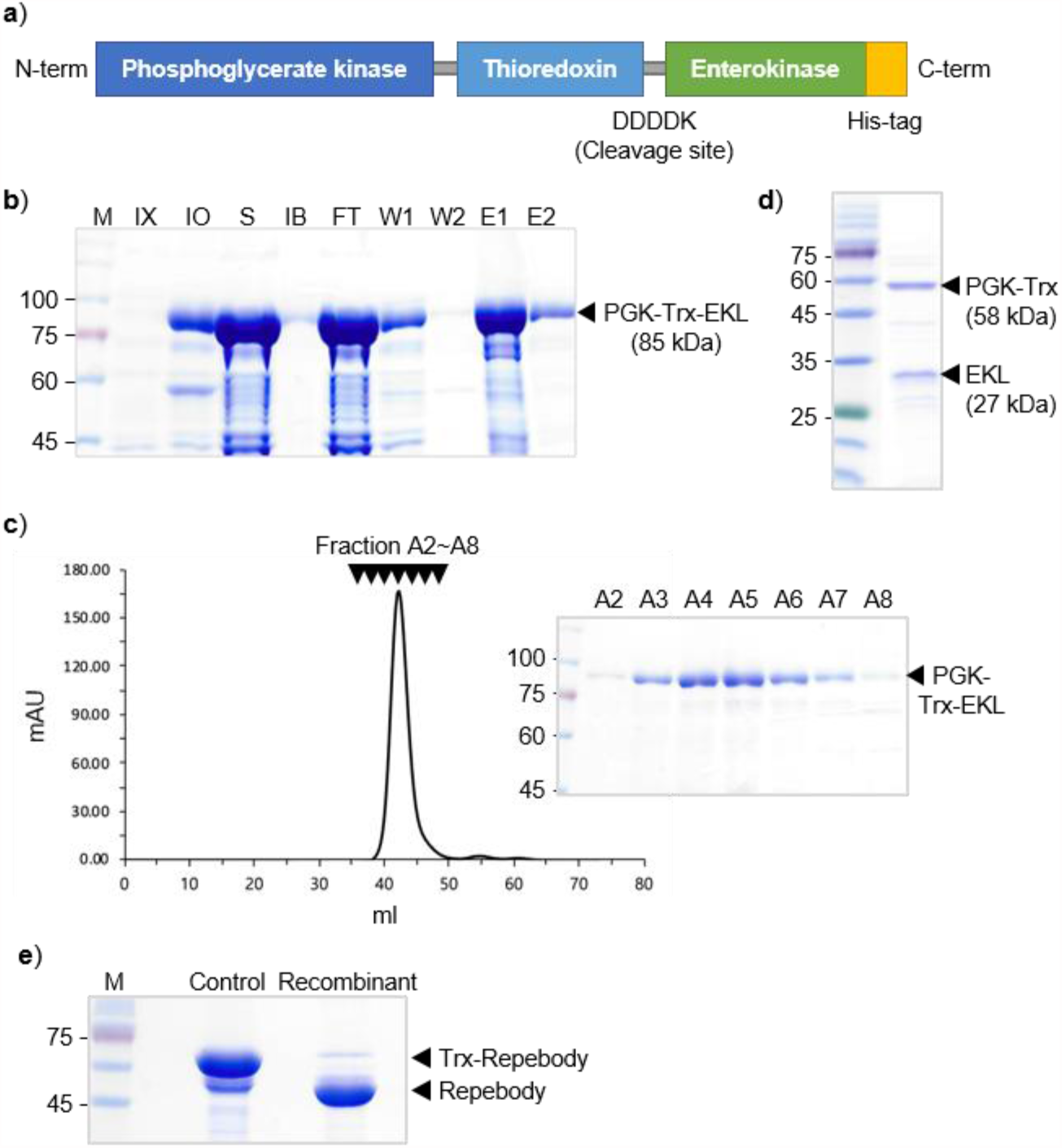
Facilitating the soluble expression of fully bioactive enterokinase through a genetic combination of phosphoglycerate kinase (PGK) and thioredoxin (Trx). **a**) Tandemly linked solubility enhancers (PGK and Trx) were genetically incorporated at the N-terminal of enterokinase (EKL). **b**) SDS-PAGE analysis for bacterial expression and purification patterns of PGK-Trx-EKL. **c**) Size-exclusion chromatography profile of PGK-Trx-EKL protein. All elution fractions were displayed through SDS-PAGE. Protein cleavage assay revealed that **d**) PGK-Trx-EKL was spontaneously cleaved, and **e**) has improved proteolytic ability compared to parent PGK-EKL.

Furthermore, we genetically engineered Trx-EKL with PGK proteins as a solubilizing tag. SDS-PAGE analysis displayed that PGK-fused Trx-EKL (PGK-Trx-EKL) was highly expressed in a soluble form, and insoluble aggregates were dramatically decreased, compared to Trx-EKL protein (**Figure 4b**). The purity of PGK-Trx-EKL was further increased by using SEC, and the resulting protein was eluted as a single peak with high purity (**Figure 4c**). Following the buffer change to an EKL reaction buffer, almost all PGK-Trx-EKL proteins were found to be spontaneously cleaved into PGK-Trx and EKL proteins (**Figure 4d**). As shown in **Figure 2d**, the proteolytic activity of PGK-tagged EKL (PGK-EKL) was relatively lower than other engineered EKL proteins. To determine if promoted disulfide bond formation by Trx tag can restore the biological function of PGK-tagged EKL, we performed the protein cleavage assay (**Figure 4e**) and found that Trx-tagged PGK-EKL protein has a higher proteolytic activity than its parent form (PGK-EKL). Based on the results, we demonstrated that each fusion protein has profound and cooperative effects on the intrinsic properties of EKL including solubility and activity. Given our findings and the fact that protein folding occurred in nature is the well-controlled biochemical process by several molecular chaperones and accessory proteins (Kim, Hipp, Bracher, Hayer-Hartl, & Hartl, 2013), the combinatorial design of solubilizing tags based on their functional complementarity could provide an alternative and practical route for cost-effective production of highly soluble and properly folded target proteins in bacteria.

## 4. Conclusion

This research presents high-level soluble expression of human enterokinase in bacteria by genetic engineering with two tandemly linked solubilizing proteins. Bacterial protein expression is highly demanded in the field of biomedical research and biotechnology to ensure cost-effective production of biological agents (Chen, 2012). Although the manipulation of bacterial expression system is relatively easy and simple, the establishment of effective methods to improve the solubility of poorly expressed proteins remains an obstacle to be addressed. Typically, the genetic fusion of protein-of-interest with a solubilizing protein is one of the most promising options to facilitate effective bacterial expression (Costa, Almeida, Castro, & Domingues, 2014; Khow & Suntrarachun, 2012). Despite advantages of solubilizing protein use, a majority of proteins are still difficult to be produced in bacteria. In fact, the ability of solubility-enhancing tags is primarily derived from their unique biological properties such as disulfide bond formation, increased hydrophilicity, and chaperone-like activity (de Marco, 2009; Lebendiker & Danieli, 2014; Raran-Kurussi & Waugh, 2012). Therefore, in order to expand the space for bacterial production of low-expressing proteins, comprehensive studies to find the most suitable combination of fusion proteins should be investigated further.

To validate this proof-of-concept, we employed three types of solubilizing tag to facilitate high-yield bacterial production of enterokinase. Among them, phosphoglycerate kinase (PGK) showed notable ability to improve the solubility of partner proteins, but the proteolytic activity of PGK-fused enterokinase was substantially decreased. This result may be because PGK is unable to promote disulfide bond formation, a prerequisite for the structural integrity and biological function of proteins. To restore its intrinsic activity, we subsequently introduced thioredoxin (Trx) to PGK-enterokinase for the effective formation of correct disulfide bridges as shown in **Scheme 1**, resulting in high-level production of soluble enterokinase with a fully retained proteolytic activity. Another Trx-fused enterokinase construct that contains maltose-binding protein (MBP) instead of PGK also showed both high solubility and bioactivity. These findings suggest that the cooperative action of fusion proteins, which plays a distinct role, can significantly increase the bacterial expression of fully active enterokinase by facilitating the protein folding process and by establishing a suitable environment to increase protein solubility. The final composition of these enterokinases using two functionalizing tags (PGK-Trx or MBP-Trx) can be used for a wide range of downstream processes, especially for production of untagged recombinant proteins on a laboratory or industrial scale. Taken together, we believe that our approach can contribute to expanding the application range of bacterial expression systems for the production of diverse biologically active and soluble recombinant proteins in a cost-effective manner.

## Acknowledgements

This research was supported by Basic Science Research Program (2019R1I1A3A01047208 and 2019R1I1A1A01058773) and Regional Leading Research Center (2020R1A5A8019180) through the National Research Foundation of Korea (NRF), and National Research Facilities & Equipment Center (2019R1A6C1010006). This study was also supported by 2018 Grant (PoINT) from Kangwon National University. All grants were funded by the Korean government (Ministry of Education and Ministry of Science and ICT).

## Author contributions

Jinhak Kwon and Hyeongjun Cho conceived the idea, designed, and constructed research materials, and performed the experiments. Yiseul Ryu and Seungmin Kim supported the experiments. Jinhak Kwon, Hyeongjun Cho and Yiseul Ryu wrote the paper. Yiseul Ryu and Joong-jae Lee edited the paper. Joong-jae Lee supervised this research. All authors analyzed data and discussed the results.

