## Supplemental Information for "A Combination Strategy of Solubility Enhancers for Effective Production of Soluble and Bioactive Human Enterokinase"

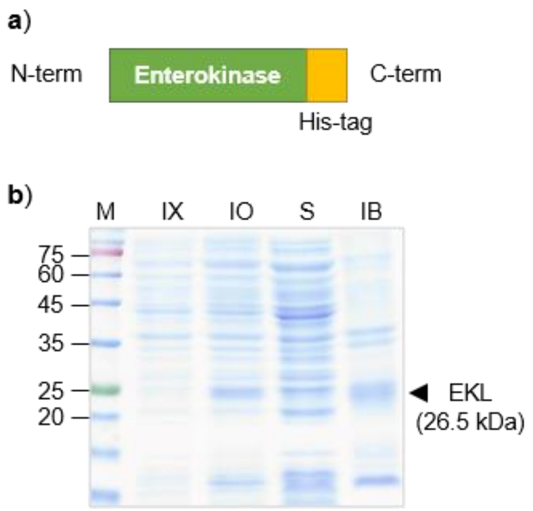


**Figure S1. Bacterial expression of human enterokinase (hEKL). a**) Human enterokinase light chain (EKL) was genetically inserted into pET21a expression vector. **b**) SDS-PAGE analysis showed that hEKL was poorly expressed with low solubility. IX: Before IPTG induction. IO: After IPTG induction. S: Soluble fraction. IB: Inclusion body.

```
        10         20         30         40         50
MGSKRGISSR HHSLSSYEIM FAALFAILVV LCAGLIAVSC LTIKESQRGA
        60         70         80         90        100
ALGQSHEARA TFKITSGVTY NPNLQDKLSV DFKVLAFDLQ QMIDEIFLSS
       110        120        130        140        150
NLKNEYKNSR VLQFENGSII VVFDLFFAQW VSDENVKEEL IQGLEANKSS
       160        170        180        190        200
QLVTFHIDLN SVDILDKLTT TSHLATPGNV SIECLPGSSP CTDALTCIKA
       210        220        230        240        250
DLFCDGEVNC PDGSDEDNKM CATVCDGRFL LTGSSGSFQA THYPKPSETS
       260        270        280        290        300
VVCQWIIRVN QGLSIKLSFD DFNTYYTDIL DIYEGVGSSK ILRASIWETN
       310        320        330        340        350
PGTIRIFSNQ VTATFLIESD ESDYVGFNAT YTAFNSSELN NYEKINCNFE
       360        370        380        390        400
DGFCFWVQDL NDDNEWERIQ GSTFSPFTGP NFDHTFGNAS GFYISTPTGP
       410        420        430        440        450
GGRQERVGLL SLPLDPTLEP ACLSFWYHMY GENVHKLSIN ISNDQNMEKT
       460        470        480        490        500
VFQKEGNYGD NWNYGQVTLN ETVKFKVAFN AFKNKILSDI ALDDISLTYG
       510        520        530        540        550
ICNGSLYPEP TLVPTPPPEL PTDCGGPFEL WEPNTTFSST NFPNSYPNLA
       560        570        580        590        600
FCVWILNAQK GKNIQLHFQE FDLENINDVV EIRDGEEADS LLLAVYTGPG
       610        620        630        640        650
PVKDVFSTTN RMTVLLITND VLARGGFKAN FTTGYHLGIP EPCKADHFQC
       660        670        680        690        700
KNGECVPLVN LCDGHLHCED GSDEADCVRF FNGTTNNNGL VRFRIQSIWH
       710        720        730        740        750
TACAENWTTQ ISNDVCQLLG LGSGNSSKPI FPTDGGPFVK LNTAPDGHLI
       760        770        780        790        800
LTPSQQCLQD SLIRLQCNHK SCGKKLAAQD ITPKIVGGSN AKEGAWPWVV
       810        820        830        840        850
GLYYGGRLLC GASLVSSDWL VSAAHCVYGR NLEPSKWTAI LGLHMKSNLT
       860        870        880        890        900
SPQTVPRLID EIVINPHYNR RRKDNDIAMM HLEFKVNYTD YIQPICLPEE
       910        920        930        940        950
NQVFPPGRNC SIAGWGTVVY QGTTANILQE ADVPLLSNER CQQQMPEYNI
       960        970        980        990       1000
TENMICAGYE EGGIDSCQGD SGGPLMCQEN NRWFLAGVTS FGYKCALPNR
      1010
PGVYARVSRF TEWIQSFLH
```

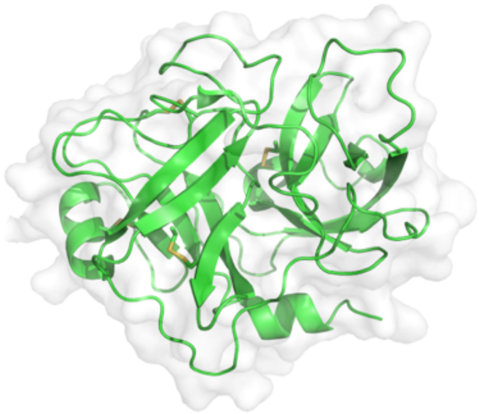

**Figure S2. Amino acid sequence and structure of the light chain of human enterokinase (hEKL).** Complete amino acid sequence of hEKL. The catalytic domain (Peptidase S1) of hEKL used in this study is highlighted in red color. Total 4 disulfide bonds are identified in three-dimensional structure of the light chain of hEKL (PDB ID 4DGJ).

**Table S1. Complete amino acid sequences of engineered enterokinases in this study.** Enterokinase protein, thioredoxin, phosphoglycerate kinase, and maltose-binding protein are colored in blue, green, purple, and orange, respectively. Enterokinase recognition sequence is highlighted in red color.

| |
|---|
| **>Enterokinase light chain (EKL)**<br>MAIVGGSNAKEGAWPWVVGLYYGGRLLCGASLVSSDWLVSAAHCVYGRNLEPSKWTAILGLHMKSNLTSPQTVPRLIDEI<br>VINPHYNRRRKDNDIAMMHLEFKVNYTDYIQPICLPEENQVFPPGRNCSIAGWGTVVYQGTTANILQEADVPLLSNERCQ<br>QQMPEYNITENMICAGYEEGGIDSCQGDSGGPLMCQENNRWFLAGVTSFGYKCALPNRPGVYARVSRFTEWIQSFLHLEH<br>HHHHH |
| **>Thioredoxin-tagged EKL (Trx-EKL)**<br>MSDKIIHLTDDSFDTDVLKADGAILVDFWAEWCGPCKMIAPILDEIADEYQGKLTVAKLNIDQNPGTAPKYGIRGIPTLL<br>LFKNGEVAATKVGALSKGQLKEFLDANLAGSGSGSSGLVPRGSGMKETAAAKFERQHMDSPDLGTDDDDKAMAIVGGSNA<br>KEGAWPWVVGLYYGGRLLCGASLVSSDWLVSAAHCVYGRNLEPSKWTAILGLHMKSNLTSPQTVPRLIDEIVINPHYNRR<br>RKDNDIAMMHLEFKVNYTDYIQPICLPEENQVFPPGRNCSIAGWGTVVYQGTTANILQEADVPLLSNERCQQQMPEYNIT<br>ENMICAGYEEGGIDSCQGDSGGPLMCQENNRWFLAGVTSFGYKCALPNRPGVYARVSRFTEWIQSFLHLEHHHHHH |
| **>Phosphoglycerate kinase-tagged EKL (PGK-EKL)**<br>MSVIKMTDLDLAGKRVFIRADLNVPVKDGKVTSDARIRASLPTIELALKQGAKVMVTSHLGRPTEGEYNEEFSLLPVVNY<br>LKDKLSNPVRLVKDYLDGVDVAEGELVVLENVRFNKGEKKDDETLSKKYAALCDVFVMDAFGTAHRAQASTHGIGKFADV<br>ACAGPLLAAELDALGKALKEPARPMVAIVGGSKVSTKLTVLDSLSKIADQLIVGGGIANTFIAAQGHDVGKSLYEADLVD<br>EAKRLLTTCNIPVPSDVRVATEFSETAPATLKSVNDVKADEQILDIGDASAQELAEILKNAKTILWNGPVGVFEFPNFRK<br>GTEIVANAIADSEAFSIAGGGDTLAAIDLFGIADKISYISTGGGAFLEFVEGKVLPAVAMLEERAKKGGGGSGGGGSGGG<br>GSIVGGSNAKEGAWPWVVGLYYGGRLLCGASLVSSDWLVSAAHCVYGRNLEPSKWTAILGLHMKSNLTSPQTVPRLIDEI<br>VINPHYNRRRKDNDIAMMHLEFKVNYTDYIQPICLPEENQVFPPGRNCSIAGWGTVVYQGTTANILQEADVPLLSNERCQ<br>QQMPEYNITENMICAGYEEGGIDSCQGDSGGPLMCQENNRWFLAGVTSFGYKCALPNRPGVYARVSRFTEWIQSFLHLEH<br>HHHHH |
| **>Maltose binding protein-tagged Trx-EKL (MBP-Trx-EKL)**<br>MKIEEGKLVIWINGDKGYNGLAEVGKKFEKDTGIKVTVEHPDKLEEKFPQVAATGDGPDIIFWAHDRFGGYAQSGLLAEI<br>TPDKAFQDKLYPFTWDAVRYNGKLIAYPIAVEALSLIYNKDLLPNPPKTWEEIPALDKELKAKGKSALMFNLQEPYFTWP<br>LIAADGGYAFKYENGKYDIKDVGVDNAGAKAGLTFLVDLIKNKHMNADTDYSIAEAAFNKGETAMTINGPWAWSNIDTSK<br>VNYGVTVLPTFKGQPSKPFVGVLSAGINAASPNKELAKEFLENYLLTDEGLEAVNKDKPLGAVALKSYEEELAKDPRIAA<br>TMENAQKGEIMPNIPQMSAFWYAVRTAVINAASGRQTVDEALKDAQTGSGGGGSGGGGSMSDKIIHLTDDSFDTDVLKAD<br>GAILVDFWAEWCGPCKMIAPILDEIADEYQGKLTVAKLNIDQNPGTAPKYGIRGIPTLLLFKNGEVAATKVGALSKGQLK<br>EFLDANLAGSGSGSSGLVPRGSGMKETAAAKFERQHMDSPDLGTDDDDKAMAIVGGSNAKEGAWPWVVGLYYGGRLLCGA<br>SLVSSDWLVSAAHCVYGRNLEPSKWTAILGLHMKSNLTSPQTVPRLIDEIVINPHYNRRRKDNDIAMMHLEFKVNYTDYI<br>QPICLPEENQVFPPGRNCSIAGWGTVVYQGTTANILQEADVPLLSNERCQQQMPEYNITENMICAGYEEGGIDSCQGDSG<br>GPLMCQENNRWFLAGVTSFGYKCALPNRPGVYARVSRFTEWIQSFLHLEHHHHHH |
| **>Phosphoglycerate kinase-tagged Trx-EKL (PGK-Trx-EKL)**<br>MSVIKMTDLDLAGKRVFIRADLNVPVKDGKVTSDARIRASLPTIELALKQGAKVMVTSHLGRPTEGEYNEEFSLLPVVNY<br>LKDKLSNPVRLVKDYLDGVDVAEGELVVLENVRFNKGEKKDDETLSKKYAALCDVFVMDAFGTAHRAQASTHGIGKFADV<br>ACAGPLLAAELDALGKALKEPARPMVAIVGGSKVSTKLTVLDSLSKIADQLIVGGGIANTFIAAQGHDVGKSLYEADLVD<br>EAKRLLTTCNIPVPSDVRVATEFSETAPATLKSVNDVKADEQILDIGDASAQELAEILKNAKTILWNGPVGVFEFPNFRK<br>GTEIVANAIADSEAFSIAGGGDTLAAIDLFGIADKISYISTGGGAFLEFVEGKVLPAVAMLEERAKKGGSGGGGSGGGGS<br>MSDKIIHLTDDSFDTDVLKADGAILVDFWAEWCGPCKMIAPILDEIADEYQGKLTVAKLNIDQNPGTAPKYGIRGIPTLL<br>LFKNGEVAATKVGALSKGQLKEFLDANLAGSGSGSSGLVPRGSGMKETAAAKFERQHMDSPDLGTDDDDKAMAIVGGSNA<br>KEGAWPWVVGLYYGGRLLCGASLVSSDWLVSAAHCVYGRNLEPSKWTAILGLHMKSNLTSPQTVPRLIDEIVINPHYNRR<br>RKDNDIAMMHLEFKVNYTDYIQPICLPEENQVFPPGRNCSIAGWGTVVYQGTTANILQEADVPLLSNERCQQQMPEYNIT<br>ENMICAGYEEGGIDSCQGDSGGPLMCQENNRWFLAGVTSFGYKCALPNRPGVYARVSRFTEWIQSFLHLEHHHHHH |
